## Supplementary Information for "Virtual-freezing fluorescence imaging flow cytometry"

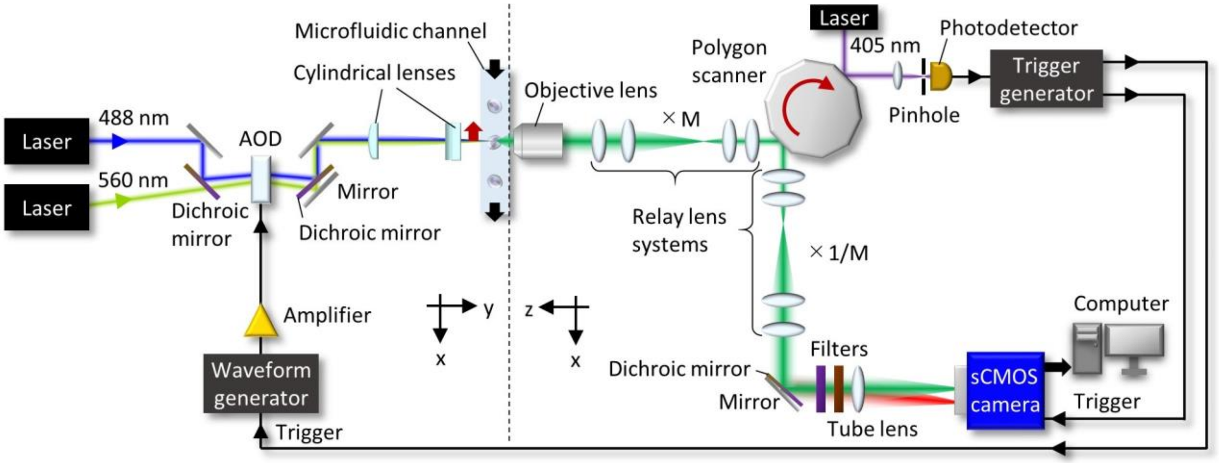


**Supplementary Fig. 1 | Complete schematic of the VIFFI flow cytometer.** The left hand side from the dashed line represents the light-sheet excitation system while the right hand side from the dashed line represents the imaging optical system and angle detection system for the polygon scanner. Note that the dimensional axes in this plane are different between the right and left hand sides.

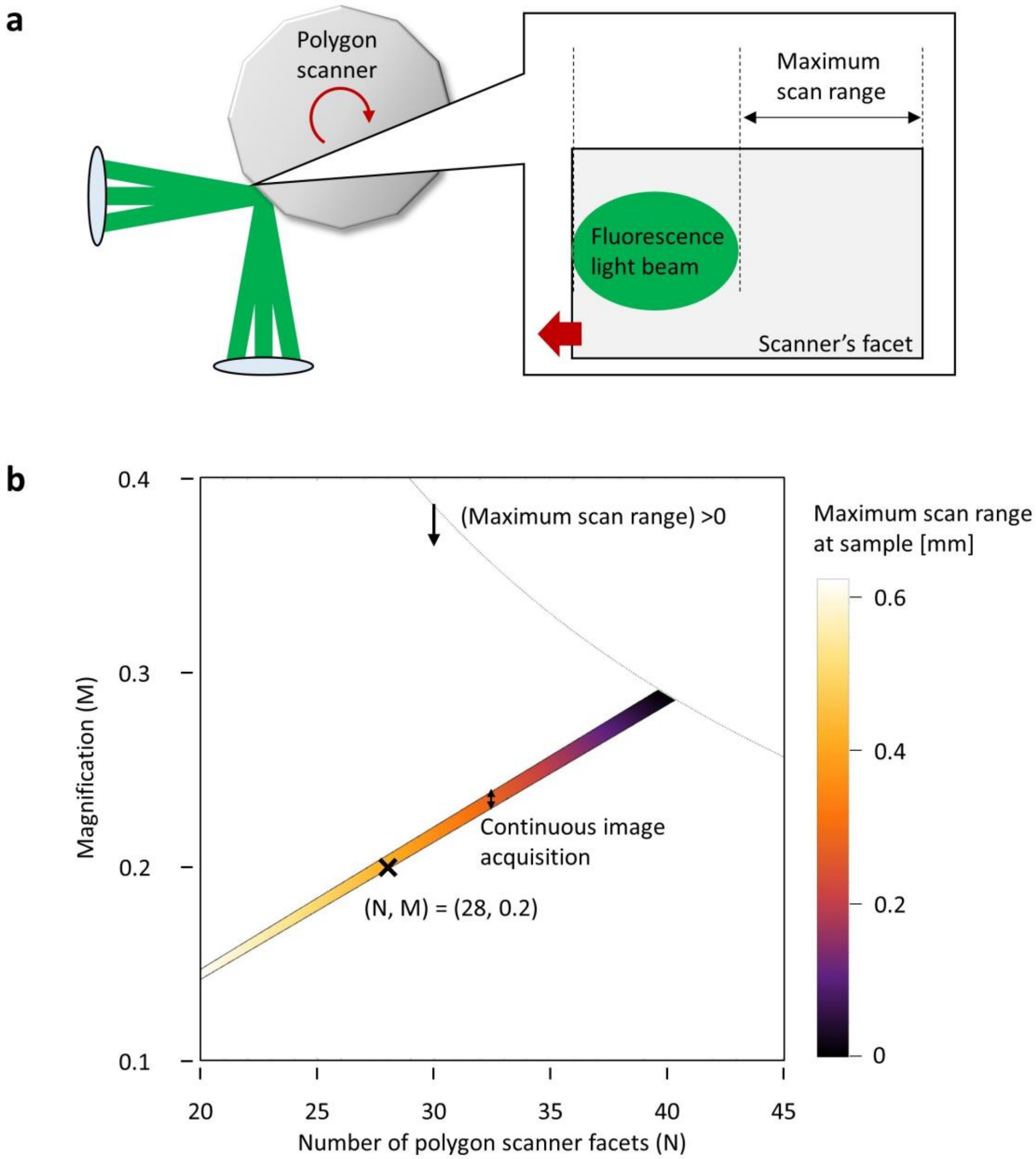


**Supplementary Fig. 2 | Design of the optical system of the VIFFI flow cytometer. a,** Configuration of the fluorescence light beam on a polygon scanner's facet. **b,** 2D plot of the maximum scan range in the object plane as a function of the number of polygon-scanner facets and the magnification of the first relay lens system.

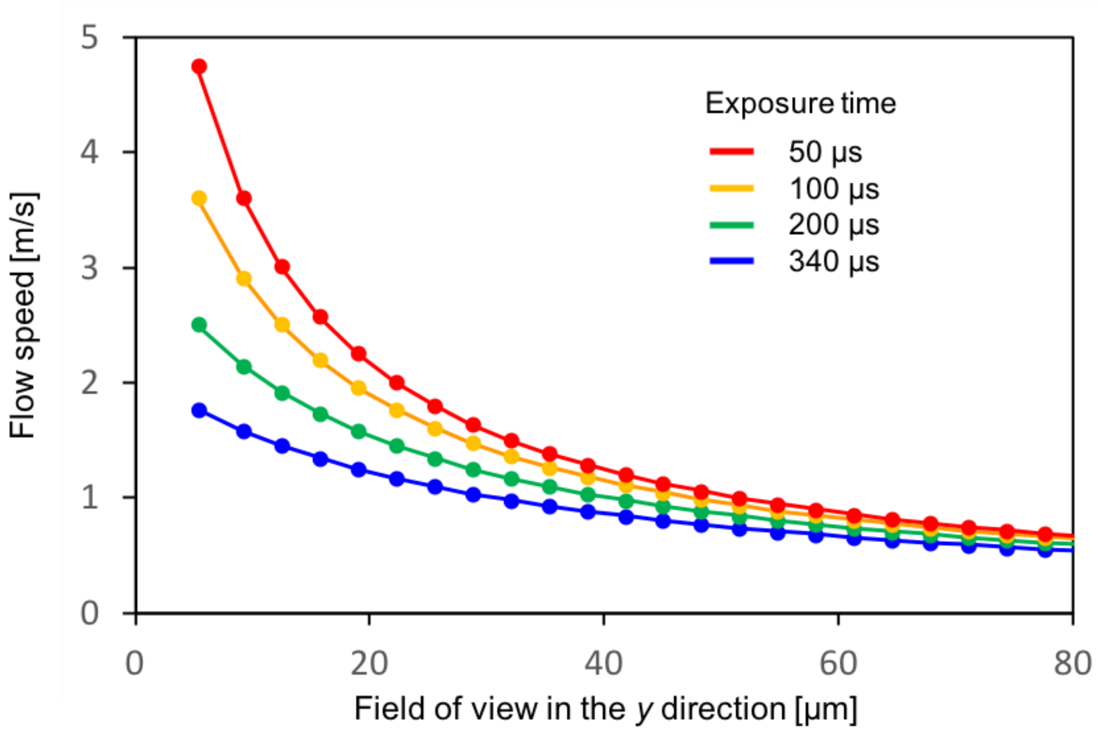


**Supplementary Fig. 3 | Relation between the cell flow speed and the FOV in the *y* direction at different camera exposure times.** The values were obtained by assuming a magnification of 20 for the imaging system and the use of the sCMOS camera model pco.edge 5.5. The cell flow speed can be increased by decreasing the FOV or shortening the camera exposure time.

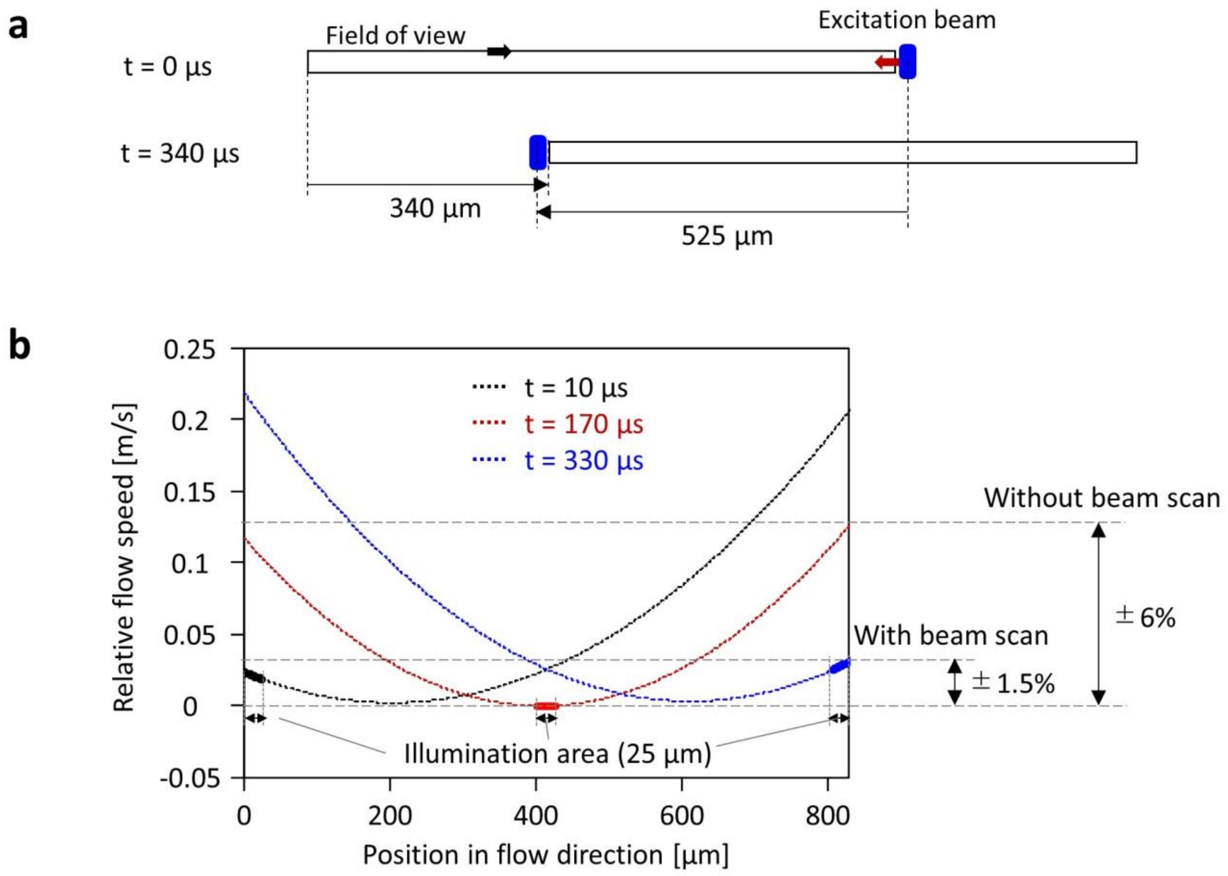


**Supplementary Fig. 4 | Details of the excitation beam scan. a,** Profile of the excitation beam scan in the object plane. **b,** Residual motion speed of flowing cells with and without the beam scan. The dotted curves represent estimated relative flow speeds at different timings. The solid curves indicate parts of the curves which correspond to illumination areas at the timings.

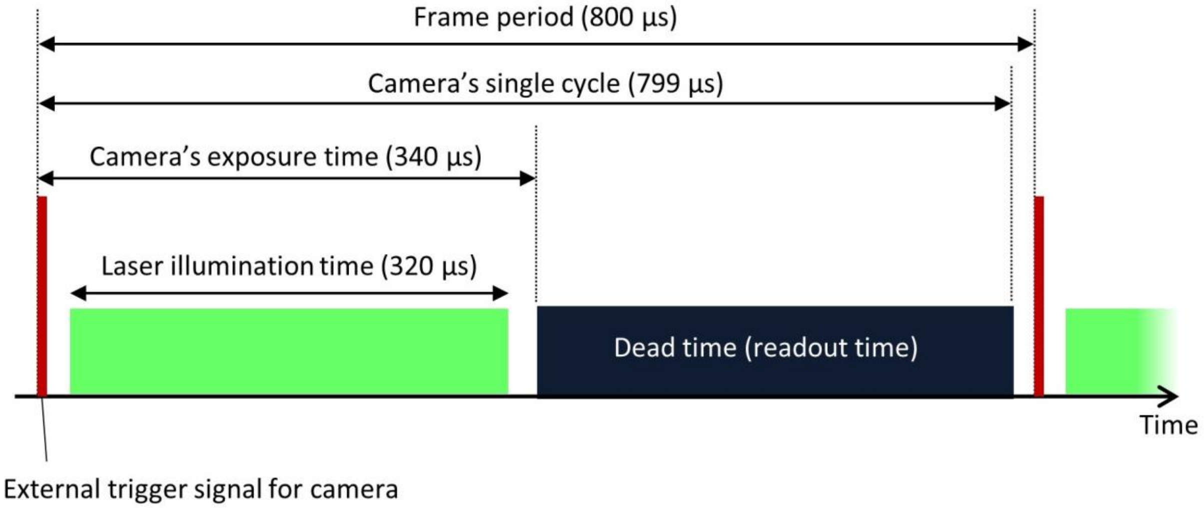


**Supplementary Fig. 5 | Schematic of the data acquisition sequence.** The sequence of the external triggering of the camera, laser illumination, and image readout repeats at a period of 800 µs. This period is slightly longer than the camera’s single cycle time (time from the external triggering to the end of readout) so that the camera is reliably triggered by every external trigger signal.

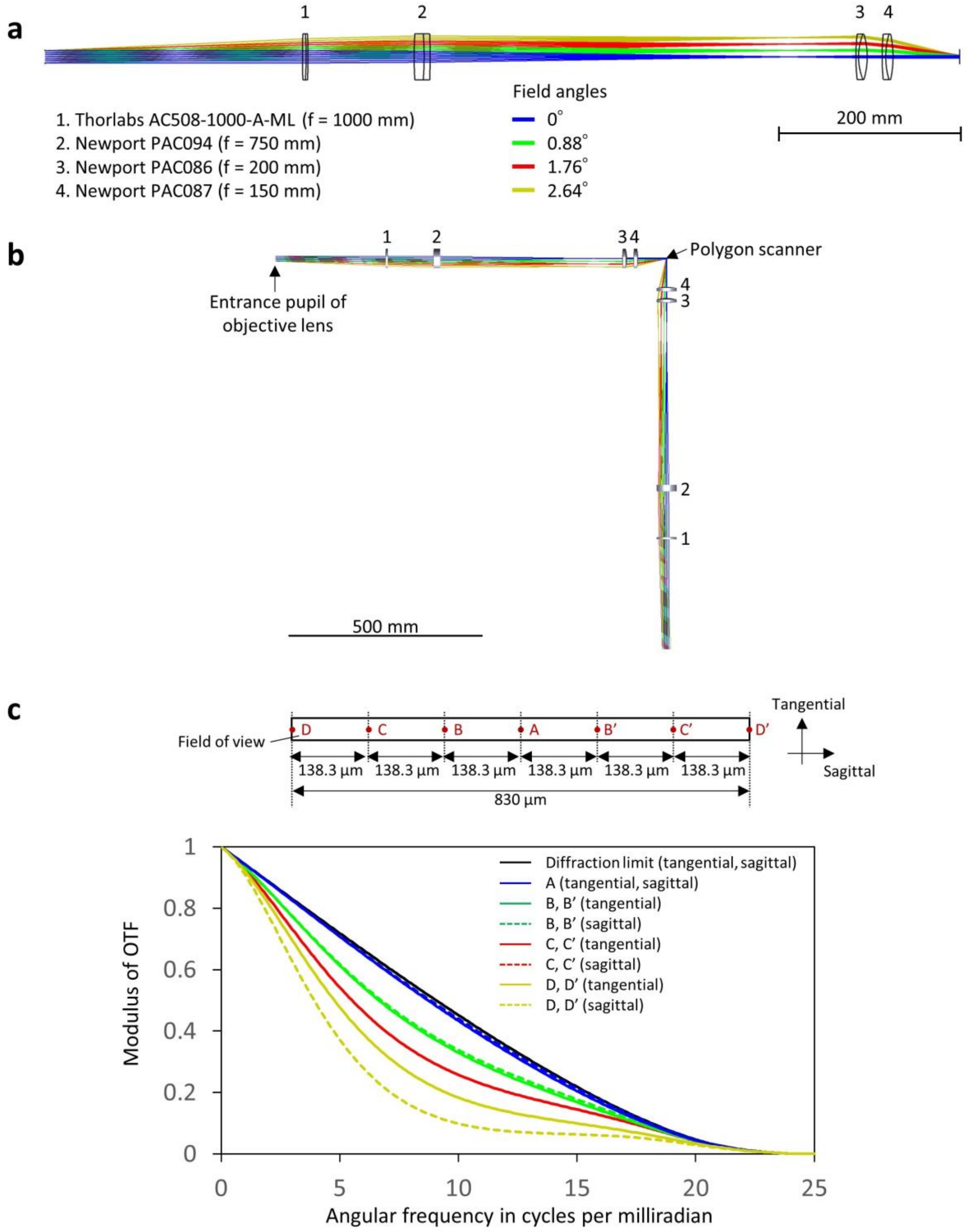


**Supplementary Fig. 6 | Design of the relay lens systems. a,** Configuration of the designed relay lens system with a magnification of 0.2. **b,** Schematic of the whole relay systems. **c,** OTFs at different positions in the FOV in the object plane.

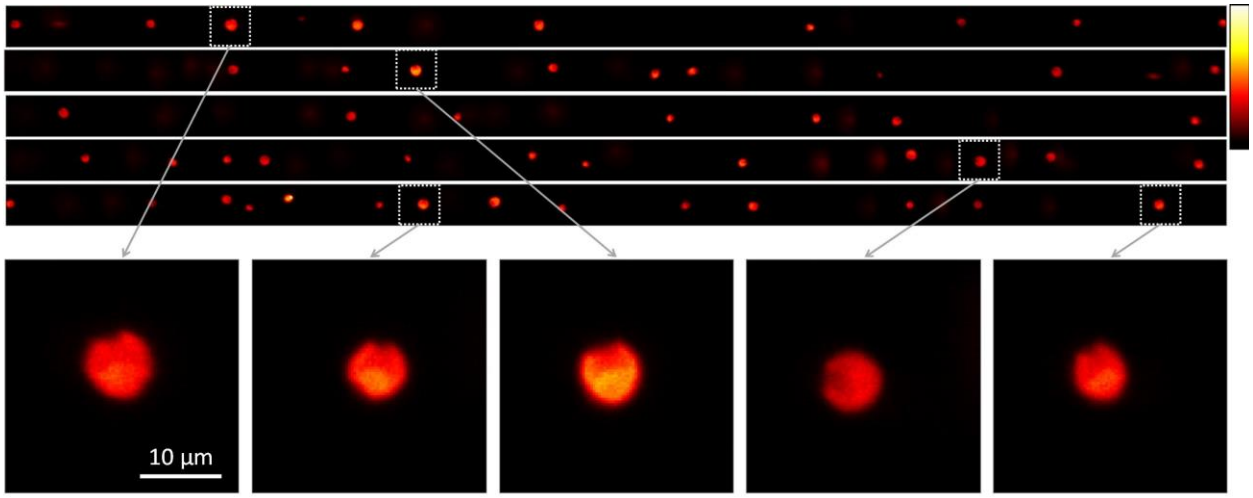


**Supplementary Fig. 7 | VIFFI flow cytometry of *Chlamydomonas reinhardtii* cells in a 1-m/s flow with subcellular resolution and a throughput of ~10,000 cells/s.** An average of ~9 autofluorescence images of *C. reinhardtii* cells were obtained in a single frame (830 μm long in the flow direction with a 30-μm overlapped length between the consecutive frames) at a flow speed of 1 m/s, demonstrating the fluorescence imaging capability with a throughput of ~10,000 cells/s. The top five images represent raw frames of the camera with a FOV of 830 μm × 28 μm while the bottom five images show zoomed parts of the frames that show characteristic morphological features (elliptical shape with an indent at the head) of *C. reinhardtii*. A laser source (Oxxius LBX-405-300-CSB-PP, 300 mW, λ = 405 nm) was used as an excitation laser.

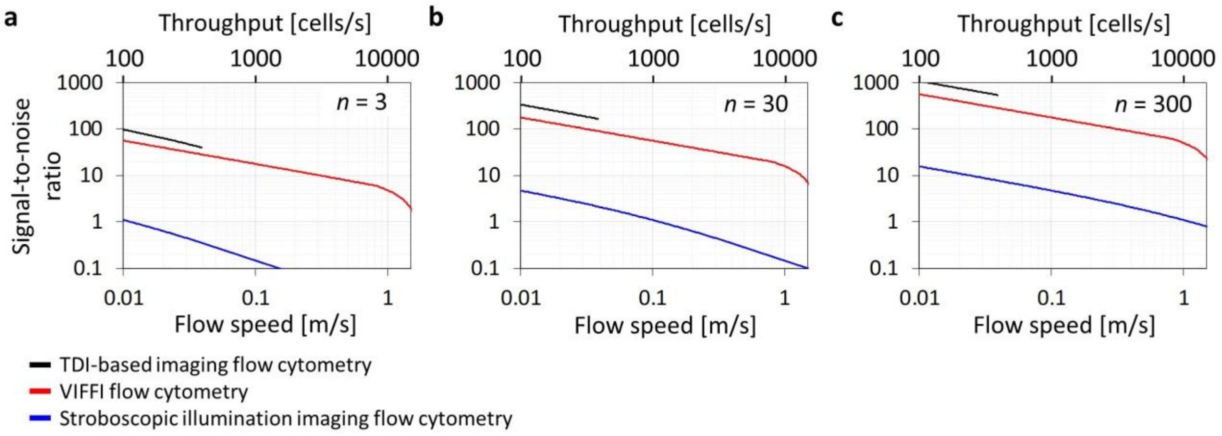


**Supplementary Fig. 8 | Estimated SNR for three different methods of IFC. a**, **b**, and **c** show the SNR at various numbers of fluorescent molecules per pixel (*n*) of 3, 30, and 300, respectively. We assume a pixel size of 0.325 μm and an average cell spacing of 100 μm. VIFFI flow cytometry has a comparable SNR with TDI-based IFC for all different numbers of fluorescent molecules per pixel. The curves of TDI-based IFC end at the flow speed of 0.04 m/s since it cannot obtain images at >0.04 m/s due to Amnis ImageStream®$^{X}$ Mark II's upper limit on the line rate. In comparison with stroboscopic illumination IFC, VIFFI flow cytometry provides a significantly higher SNR by more than a factor of 30, enabling image acquisition with a reasonable SNR at a high speed of ~1 m/s.

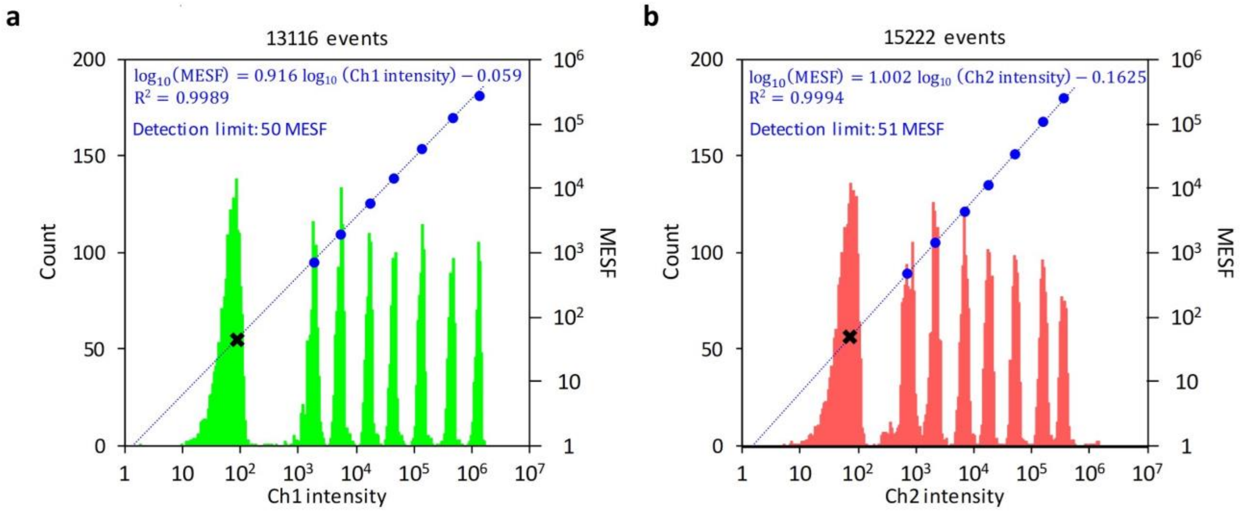


**Supplementary Fig. 9 | Evaluation of the detection sensitivity of the VIFFI flow cytometer using standard 8-peak fluorescent calibration particles. a** and **b** show results for the green channel (ch1, 510 nm – 580 nm) and the red channel (ch2, > 580 nm), respectively. The histograms show distributions of the signal intensities of the particles with different fluorescence intensities (left vertical axis). We used SPHEROTM$^{TM}$ Rainbow Calibration Particles (Spherotech Inc., catalog no. RCP-30-5A, lot no. AK02). Also, we used the microfluidic platform shown in Ref. 26 to detect non-fluorescent particles using the speed meter setup. The signal intensities were evaluated as the integrated pixel intensities in circular image areas with a 3.6-μm diameter around the center of the particles (diameter: 3 μm). Defocused images of the particles were excluded from the analysis. The blue dots indicate the average signal intensities and MESF values of the populations (right vertical axis), with regression lines and fit functions shown in blue.

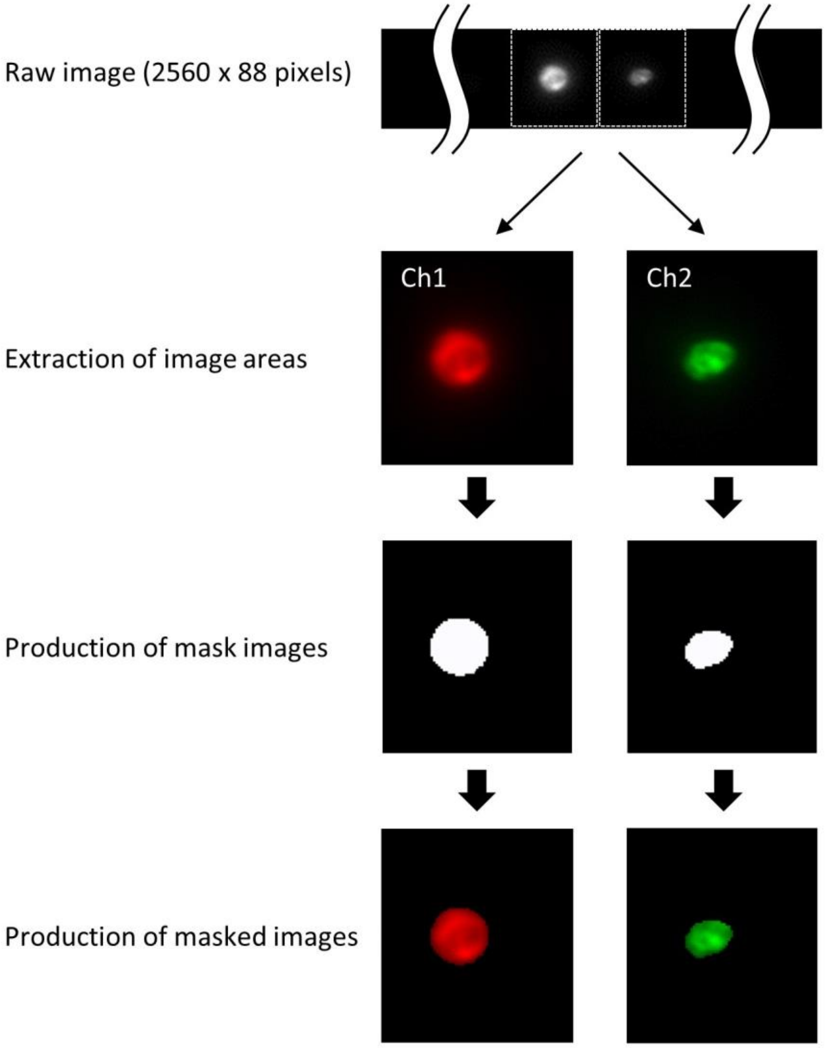


**Supplementary Fig. 10 | Flowchart for digital image processing.** Two images of a cell that correspond to different color channels are extracted from a raw image. Mask images are created from the extracted image areas, which are used for producing masked cell images.

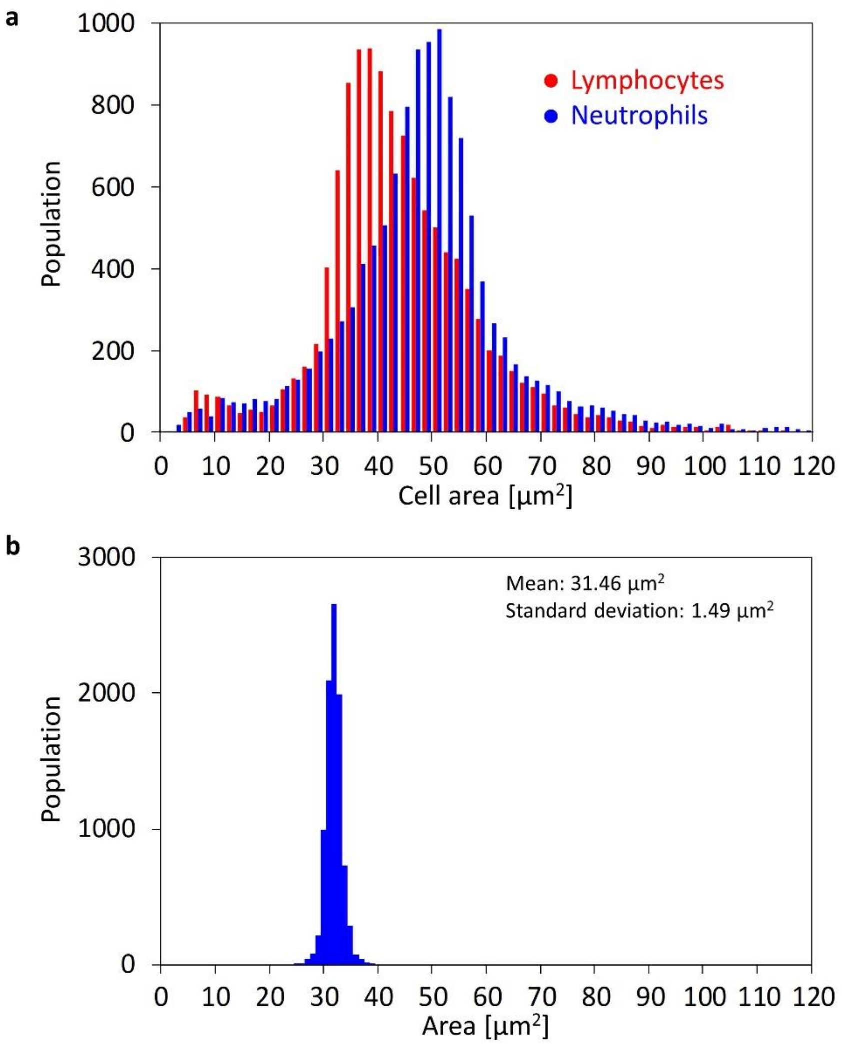


**Supplementary Fig. 11 | Statistical analysis of cell areas. a,** Histograms of murine lymphocytes and neutrophils in terms of cell area. **b,** Histogram of 6-μm fluorescent particles in terms of particle area. The narrow distribution of the histogram indicates that the distributions shown in panel **a** reflect those of the cell areas of lymphocytes and neutrophils.

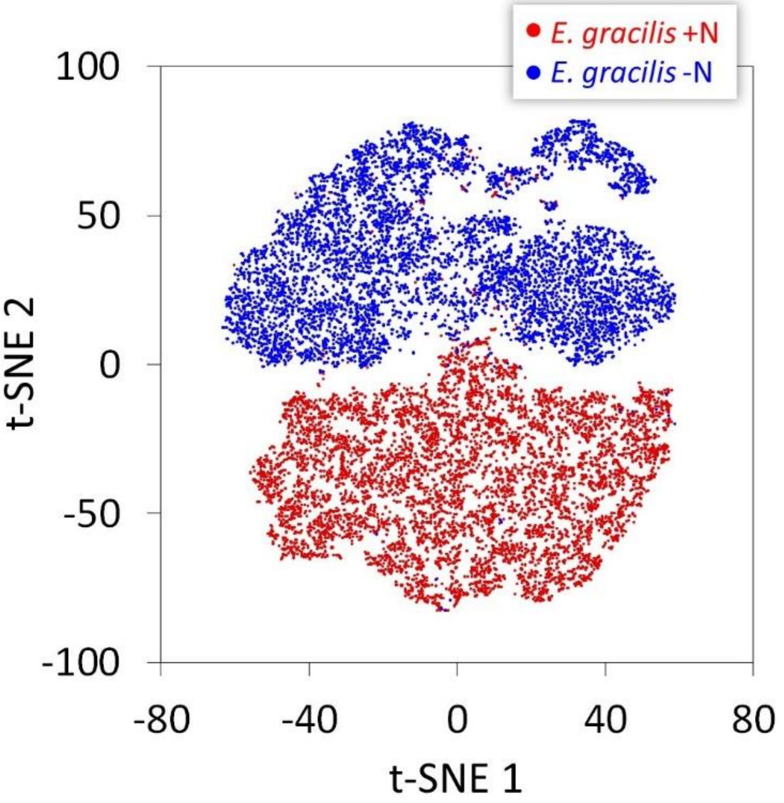


**Supplementary Fig. 12 | t-SNE plot of *E. gracilis* cells.** Red and blue dots represent *E. gracilis* +N and –N, respectively. t-SNE 1 and t-SNE 2 are obtained with 4,096 features by VGG-16.

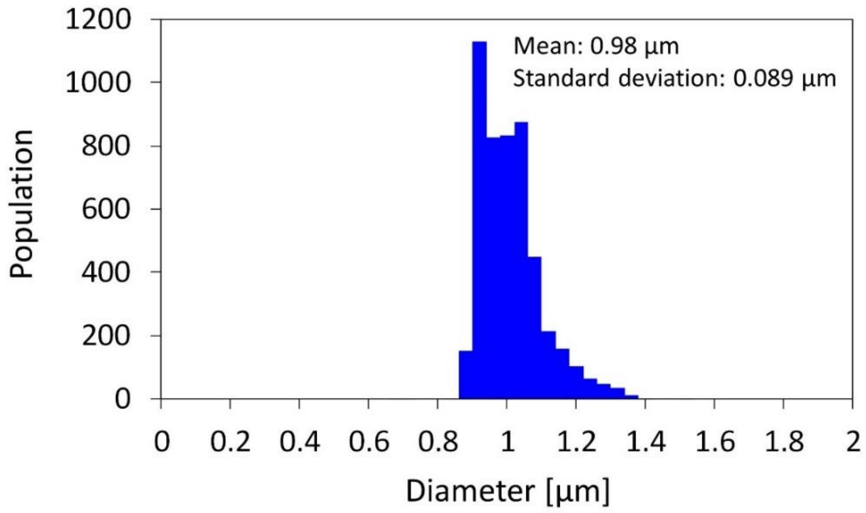


**Supplementary Fig. 13 | Histogram of 1-µm fluorescent particles in terms of particle diameter.** We evaluated the diameter by FW$e^{-1}$M of the fitted Gaussian function for the line profile in the *y* direction. The tailed shape of the distribution indicates out-of-focus particles.

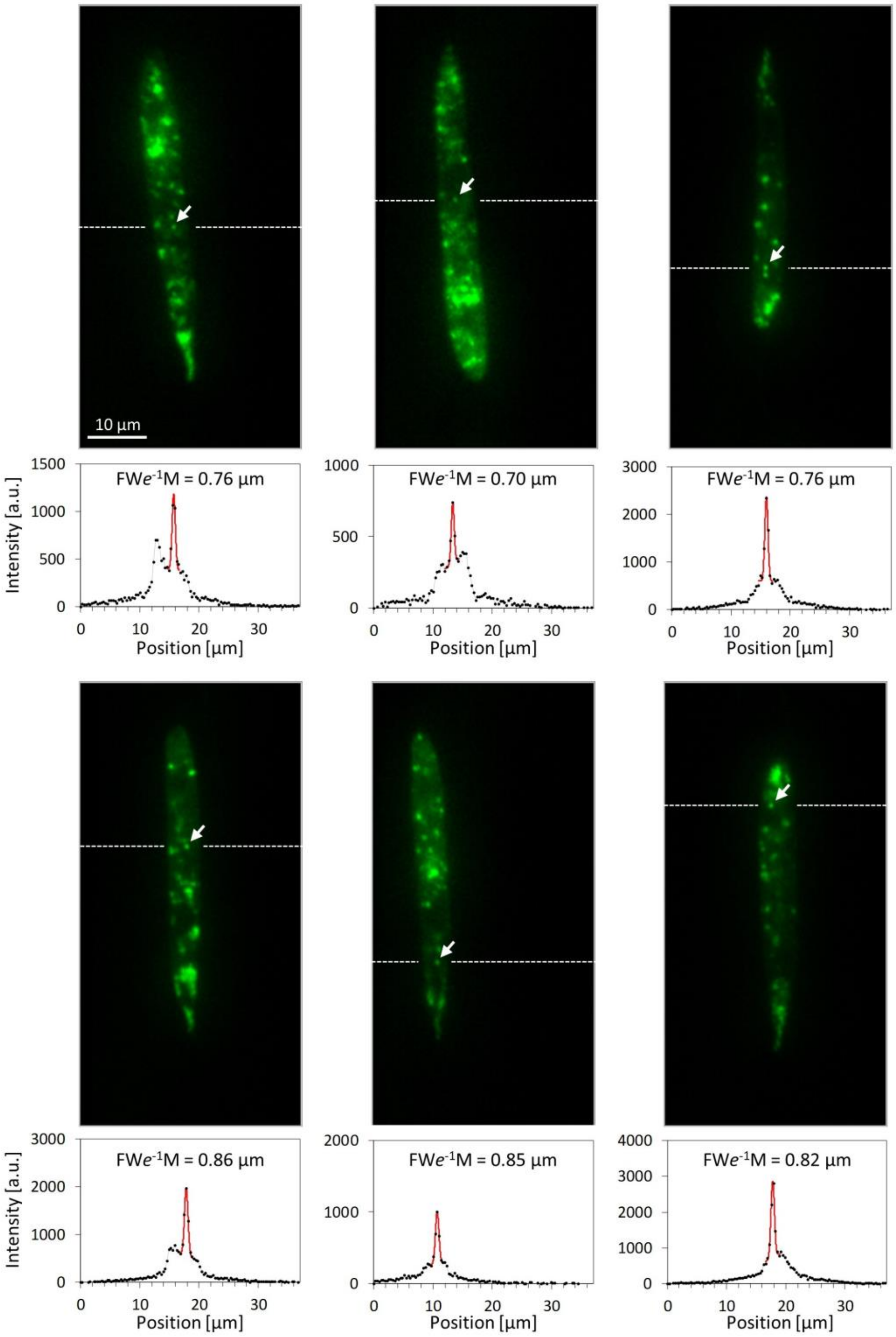


**Supplementary Fig. 14 | Evaluation of the size of small lipid droplets in *E. gracilis* cells.** The images show lipid droplets in *E. gracilis* cells. The dashed lines correspond to the horizontal axes of the plots below the images. The arrows in the images indicate lipid droplets whose diameters are evaluated in the plots. The black dots connected by the gray lines indicate pixel values. The red curves indicate fitting results for the peaks of small droplets with a Gaussian function with a slanted (linearly varying) background.

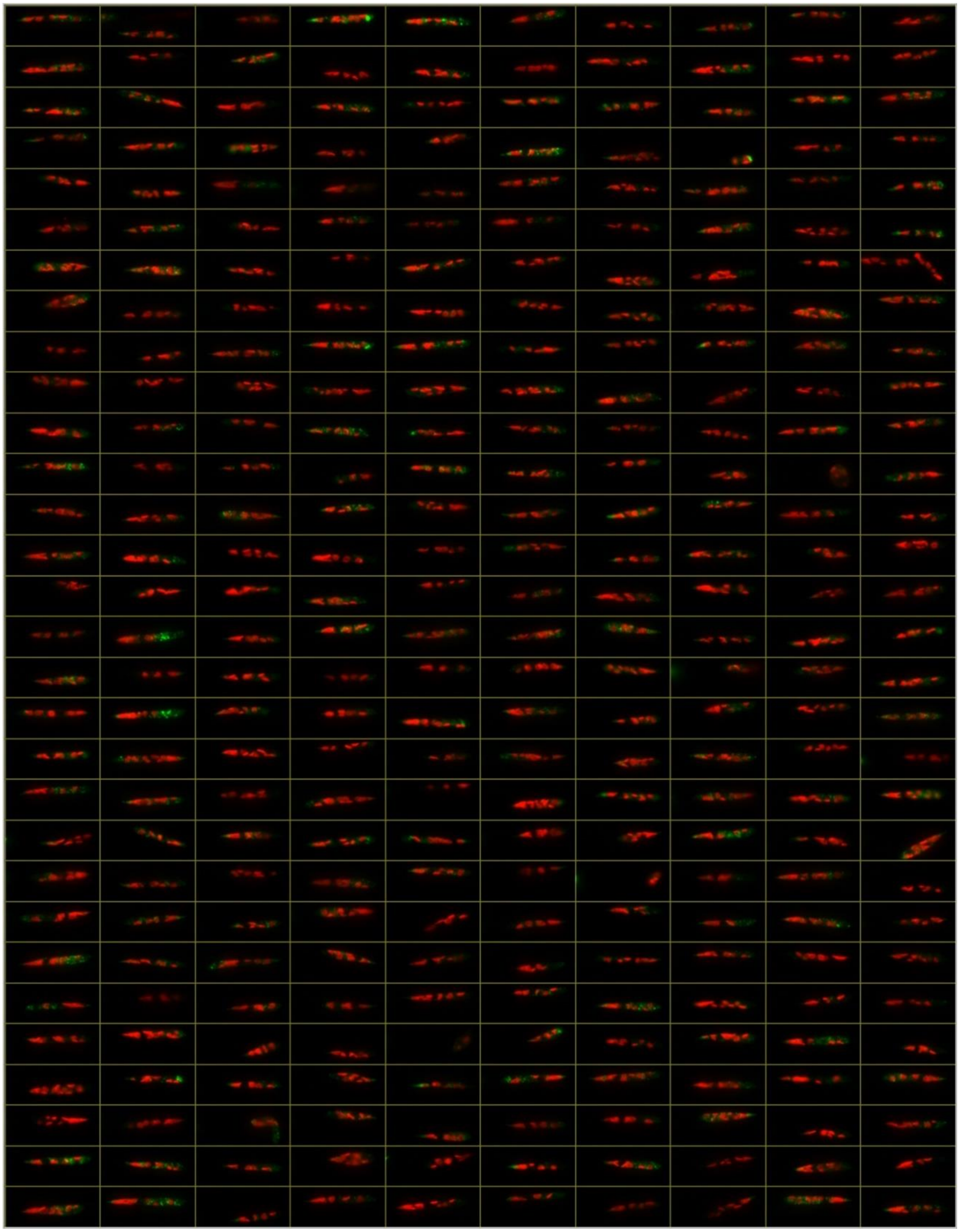

**Supplementary Fig. 15 | Images of *E. gracilis* +N cells obtained by the VIFFI flow cytometer.** The lipids (shown in green) were stained by BODIPY505/515 while chlorophyll (shown in red) was autofluorescent.

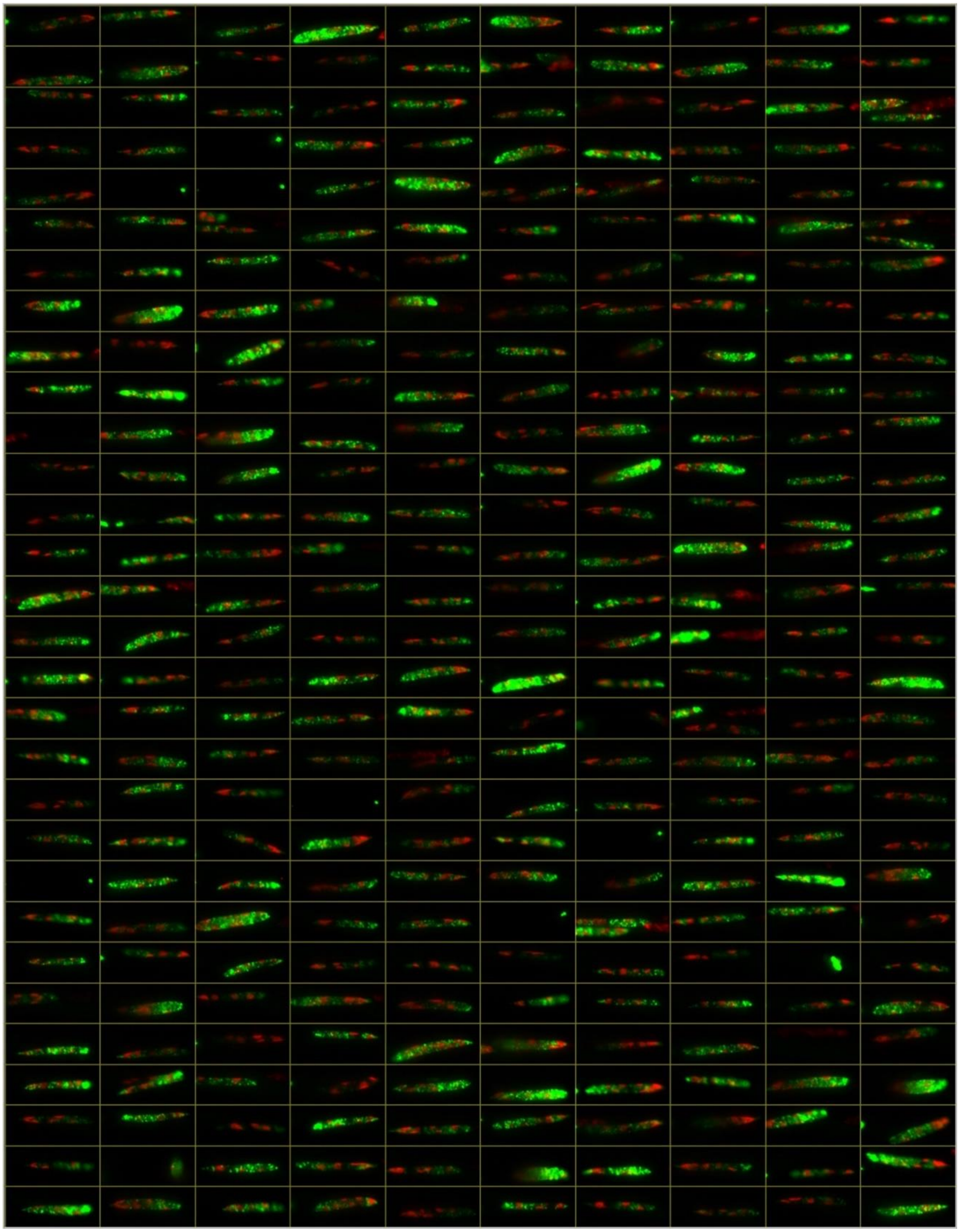

**Supplementary Fig. 16 | Images of *E. gracilis* –N cells obtained by the VIFFI flow cytometer.** The lipids (shown in green) were stained by BODIPY505/515 while chlorophyll (shown in red) was autofluorescent.

green).

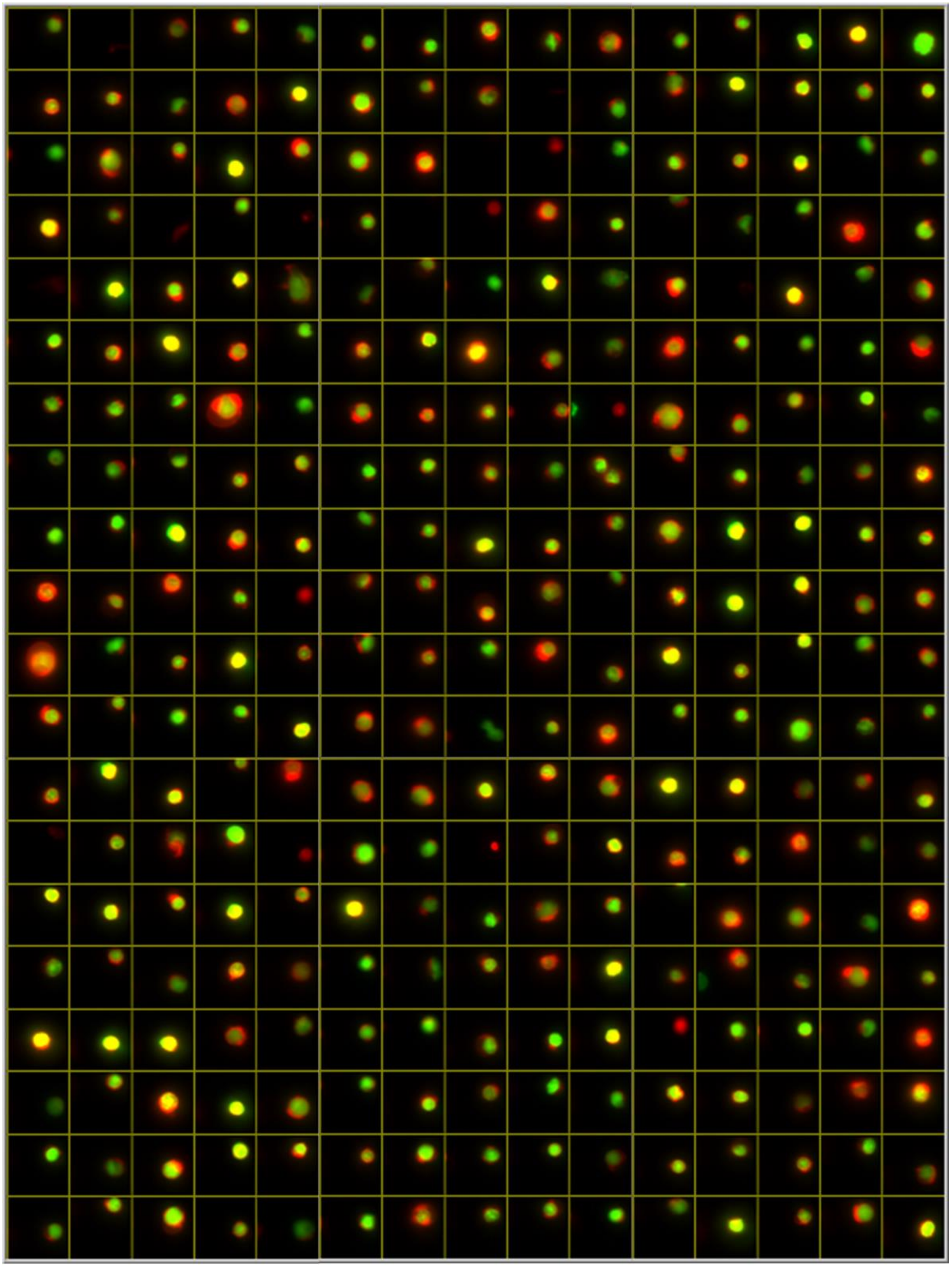

**Supplementary Fig. 17 | Images of murine lymphocytes obtained by the VIFFI flow cytometer.** The cytoplasm was stained by CellTracker Red (shown in red) while the nucleus was stained by SYTO16 (shown in green).

cytoplasm was stained by CellTracker Red (shown in red) while the nucleus was stained by SYTO16 (shown in green).

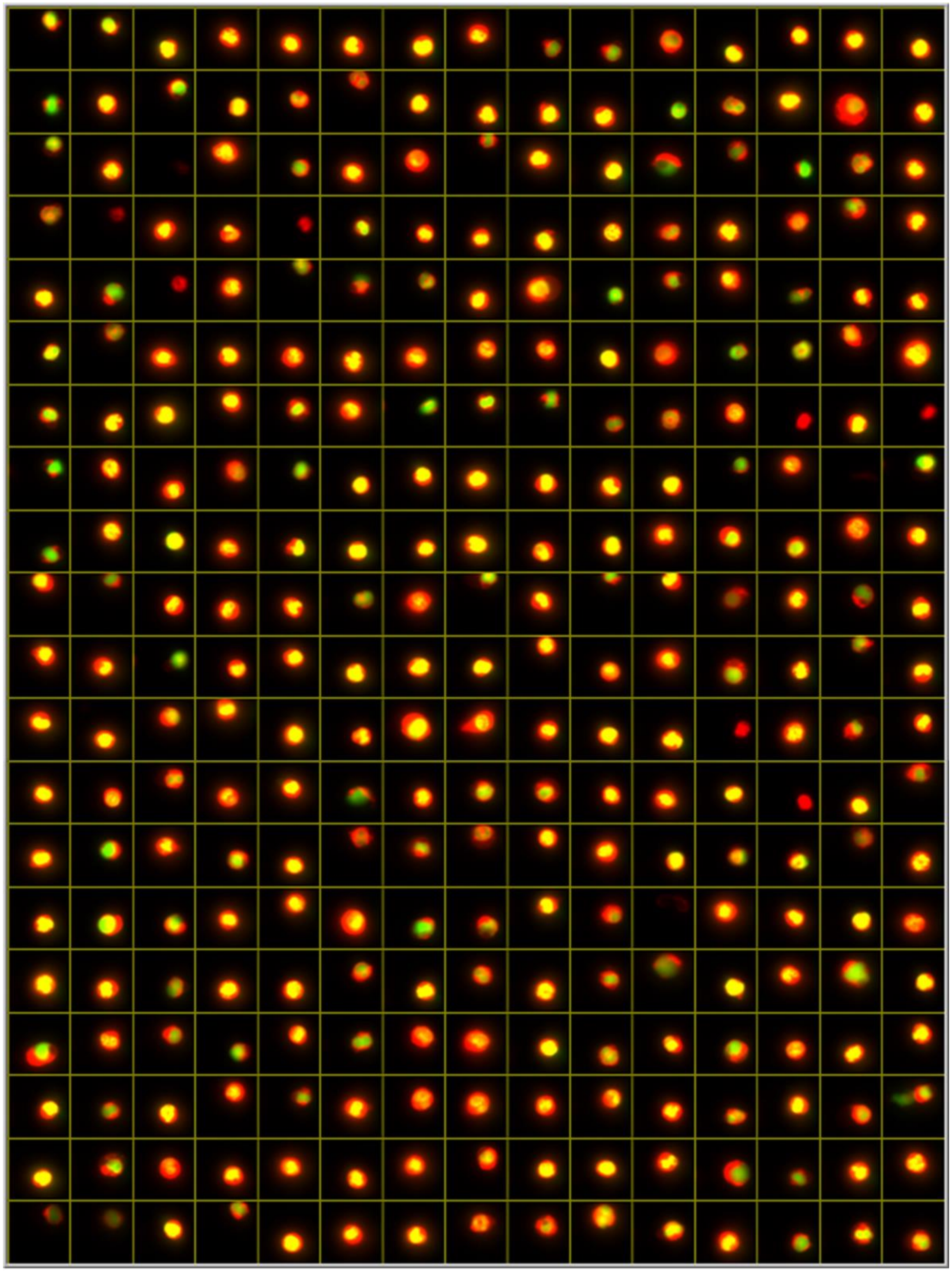

**Supplementary Fig. 18 | Images of murine neutrophils obtained by the VIFFI flow cytometer.** The cytoplasm was stained by CellTracker Red (shown in red) while the nucleus was stained by SYTO16 (shown in green).

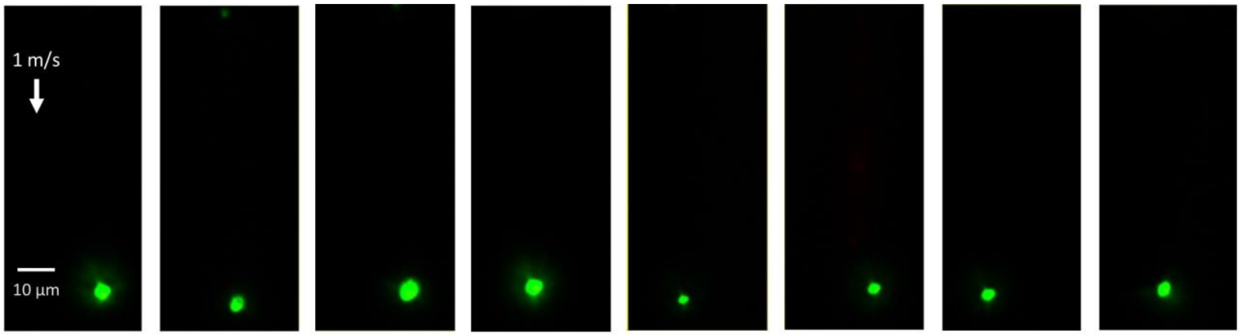


**Supplementary Fig. 19 | Images of isolated lipid droplets obtained by the VIFFI flow cytometer.** These images correspond to the near-origin peak of the *E. gracilis* –N histogram in Fig. 4c.

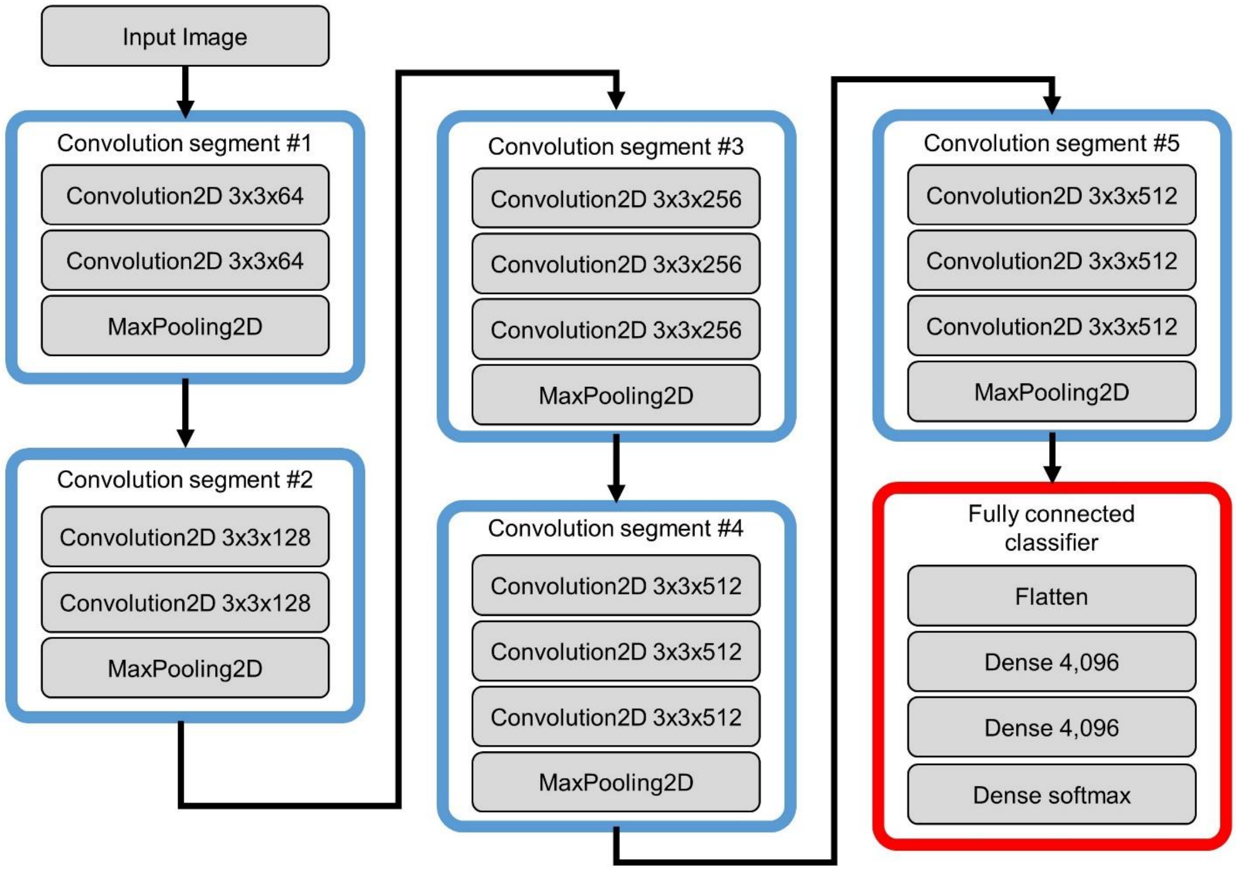


**Supplementary Fig. 20 | Network architecture of VGG16.** 4,096 features are extracted from a (multi-color) input image through five convolution segments and a fully connected classifier. Each convolution segment consists of two or three convolutional layers for feature extraction and a maximum pooling layer for reduction of data volume. The fully connected classifier downgrades the 4,096 features to one dimension to provide a classification result.

**Supplementary Table 1 | Parameter settings for the estimation of the SNRs.**

| | | VIFFI flow cytometry | | TDI-based IFC | | Stroboscopic illumination IFC | |
|---|---|---|---|---|---|---|---|
| Parameter | Symbol | Value | Note | Value | Note | Value | Note |
| Excitation beam power [mW] | $P$ | 180 | Accounting for ~10% loss at the beam scanner | 200 | Maximum output power of the 488-nm laser in ImageStream®X Mark II | 200 | Same as the left for fair comparison |
| Excitation beam diameter in the depth direction [μm] | $D_z$ | 4 | Current setting | 4 | Same as the left value for fair comparison | 4 | Same as the left value for fair comparison |
| Excitation beam diameter in the flow direction [μm] | $D_x$ | 26 | Current setting | 26 | Same as the left value for fair comparison | 666 | Full FOV in the flow direction at 20x |
| Cross section of the excitation beam [μm²] | $C$ | 104 | Approximated by $\pi D_x D_z/4$ | 104 | Approximated by $\pi D_x D_z/4$ | 104 | Approximated by $\pi D_x D_z/4$ |
| Time of the excitation beam illumination | $T$ | (vari-able) | $\min(t_{exp1}, t_{exp2}) D_x/FOV_x$ (see Eqs. S5 and S6) | (vari-able) | $D_x$/(flow speed) | (vari-able) | (pixel size)/(flow speed) |
| Absorption cross section [cm²] | $s$ | $1.16 \times 10^{-16}$ | Fluorescein | $1.16 \times 10^{-16}$ | Fluorescein | $1.16 \times 10^{-16}$ | Fluorescein |
| Quantum yield | $\eta_{yield}$ | 0.8 | Fluorescein | 0.8 | Fluorescein | 0.8 | Fluorescein |
| Photon collection efficiency of the imaging system | $\eta_{opt}$ | 0.0277 | Calibrated value of the present setup at NA = 0.75 | 0.0305 | 10% higher than the left value, NA = 0.75 | 0.0129 | Same as the left at the same NA, NA = 0.5 |
| Quantum efficiency of the image sensor | $\eta_{sensor}$ | 0.55 | Specification of pco.edge 5.5 (λ = 530 nm) | 0.8 | Specification of Hamamatsu C10000-801 (λ = 530 nm) | 0.83 | Specification of Hamamatsu ORCA-Flash 4.0 V3 (λ = 530 nm) |
| Readout noise [$e^-$ rms] | $\sigma$ | 1.7 | Specification of pco.edge 5.5 (λ = 530 nm) | 50 | Specification of Hamamatsu C10000-801 (λ = 530 nm) | 1.6 | Specification of Hamamatsu ORCA-Flash 4.0 V3 (λ = 530 nm) |

**Supplementary Table 2 | Comparison of parameters related to data acquisition speed.**

| | VIFFI flow cytometry | ImageStream®$^{X}$ Mark II |
|---|---|---|
| Pixel size | 0.325 μm | 0.33 μm[1)] |
| Flow speed | 1 m/s | 0.04 m/s[2)] |
| Effective line rate | 3.1 MHz | 0.12 MHz |
| Cell throughput at an average cell spacing of 100 μm | 10,000 cells/s | 400 cells/s |

1) Specification of ImageStream®$^{X}$ Mark II at 60-X magnification

2) INSPIRE® (ImageStreamX® System Software) User's Manual Version Mark II
